# Colour vision models: a practical guide, some simulations, and *colourvision* R package

**DOI:** 10.1101/103754

**Authors:** Felipe M. Gawryszewski

## Abstract

- Human colour vision differs from the vision of other animals. The most obvious differences are the number and type of photoreceptors in the retina. E.g., while humans are insensitive to ultraviolet (UV) light, most non-mammal vertebrates and insects have a colour vision that spans into the UV. The development of colour vision models allowed appraisals of colour vision independent of the human experience. These models are now widespread in ecology and evolution fields. Here I present a guide to colour vision modelling, run a series of simulations, and provide a R package – `colourvision` – to facilitate the use of colour vision models.
- I present the mathematical steps for calculation of the most commonly used colour vision models: Chittka (1992) colour hexagon, Endler & Mielke (2005) model, and Vorobyev & Osorio (1998) linear and log-linear receptor noise limited models (RNL). These models are then tested using identical simulated and real data. These comprise of reflectance spectra generated by a logistic function against an achromatic background, achromatic reflectance against an achromatic background, achromatic reflectance against a chromatic background, and real flower reflectance data against a natural background reflectance.
- When the specific requirements of each model are met, between model results are, overall, qualitatively and quantitatively similar. However, under many common scenarios of colour measurements, models may generate spurious values and/or considerably different predictions. Models that log-transform data and use relative photoreceptor outputs are prone to generate unrealistic results when the stimulus photon catch is smaller than the background photon catch. Moreover, models may generate unrealistic results when the background is chromatic (e.g. leaf reflectance) and the stimulus is an achromatic low reflectance spectrum.
- Colour vision models are a valuable tool in several ecology and evolution subfields. Nonetheless, knowledge of model assumptions, careful analysis of model outputs, and basic knowledge of calculation behind each model are crucial for appropriate model application, and generation of meaningful and reproducible results. Other aspects of vision not incorporated into these models should be considered when drawing conclusion from model results.

## Introduction

Animals respond to their surroundings via processing of data acquired by sensory organs (Stevens 2013). The senses can evolve in response to selective pressures from the environment as the senses can exert selective pressures into other organism morphology and behaviour. E.g. the peak of sensitivity of marine mammal photoreceptors correlates to the environmental light conditions (Fasick & Robinson 2000), and flower colours parameters may have evolved in response to the visual abilities of pollinators (Chittka & Menzel 1992; Dyer *et al*. 2012).

There are several differences in the vision of animals – between and sometimes within species – such as density and distribution of receptors in the retina, visual acuity, and presence of oil-droplets in photoreceptors cells (Cronin et al., 2014). In terms of colour vision, the most obvious differences are the type of photoreceptors present in the retina (Kelber *et al*. 2003; Osorio & Vorobyev 2008). Old world primates, including humans, are trichromats (have three cones types), with sensitivity peaks in blue, green and red regions of the light spectrum, whereas other mammals are usually dichromats (Kelber *et al*. 2003; Osorio & Vorobyev 2008; Jacobs 2009).

Most non-mammal vertebrates are tetrachromats, most insects are trichromats, and both have a colour perception that spans into the ultraviolet (Bowmaker 1998; Briscoe & Chittka 2001; Osorio & Vorobyev 2008). A fascinating illustration of how photoreceptor sensitivity may affect colour perception comes from human subjects that had gone through cataract treatment. The sensitivity curve of human blue photoreceptor actually spans into the ultraviolet (UV), but humans are UV-insensitive because pigments in the eye crystalline filters-out wavelengths below 400nm. Cataract surgery occasionally replaces the crystalline with an UV-transmitting lens, and those individuals are suddenly able to see the world differently: new patterns appear in flower petals, some garments originally perceived as black become purple, and black light are turned into blue light (Stark & Tan 1982; Cornell 2011).

Therefore, human colour perception is, in most cases, only a crude approximation of other species colour vision. Thus, studies of animal colouration can clearly benefit from appraisals of how colour patches are perceived by non-human observers. Moreover, the same colour patch may be perceived differently not only depending on the observer, but also on the context that this colour patch is exposed (e.g. background colour and environmental light conditions; (Endler 1978).

Colour vision models where firstly developed in an attempt to understand the proximate causes of human colour vision, and emulate some of human visual perceptual phenomena (Kemp *et al*. 2015). More recently, with the advent of affordable spectrometers for reflectance measurements, application of colour vision models became common place in ecology and evolution subfields. Together, some of the most important colour vision papers have been cited over 2800 times (Endler 1990 (919); Vorobyev & Osorio 1998 (601), Vorobyev *et al*. 1998 (460); Chittka 1992 (324); Chittka *et al*. 1992 (128); Endler & Mielke 2005 (445); Google Scholar search on October 31th 2016). A few examples of use of colour vision models include studies of plant-pollinator interactions (Whitney *et al*. 2009), evolution of avian plumage (Stoddard & Prum 2008), sexual selection (Amy *et al*. 2008), visual prey lures (Heiling *et al*. 2003), speciation (Carleton *et al*. 2005), mimicry (Stoddard & Stevens 2011), camouflage (Thery & Casas 2002) and aposematism (Siddiqi 2004).

As any model, colour vision models are based on certain assumptions. Knowledge of model strength and limitations are crucial to assure reproducible and meaningful results from model applications. Thus, the motivation of this paper is twofold: first to facilitate the use of colour vision models by evolutionary biologists and ecologists; secondly, to compare the consistency of between model results and how they behave in common scenarios of colour measurements. I did not aim to give an in depth analysis of the physiology of colour vision, but to provide a practical guide to the use of colour vision models, and demonstrate their limitations and strengths. Guidance on other aspects of colour vision models can be found elsewhere (Kelber *et al*. 2003; Endler & Mielke 2005; Osorio & Vorobyev 2008; Kemp *et al*. 2015; Renoult *et al*. 2017). I begin with a mathematical description of the steps for calculation of the most common colour vision models used in ecology and evolution; then I run a series of simulations using colour vision models. Both the description and the simulations serve as presentation of the accompanying R package `colourvision`.

### Colour vision models

In general, colour vision is achieved by neural opponency mechanisms (Kelber *et al*. 2003; Kemp *et al*. 2015), although exceptions to this rule do exist (Thoen *et al*. 2014). In humans, two colour opponency mechanisms appear to dominate: yellow-blue and red-green opponency channels (Kelber *et al*. 2003). Although colour vision in most other animals studied so far also seem to be based on opponency mechanisms, the exact opponency channels are usually not known (Kelber *et al*. 2003; Kemp *et al*. 2015). Nonetheless, empirical studies suggest that the exact opponency channels do not need to be known for a good prediction of behavioural responses by colour vision models (Chittka *et al*. 1992; Vorobyev & Osorio 1998; Spaethe *et al*. 2001; Cazetta *et al*. 2009).

Here I present and test four generalist colour vision models used in ecology and evolution: Chittka (1992) colour hexagon, Endler & Mielke (2005) model, and linear and log-linear versions of the Receptor Noise Limited model (Vorobyev & Osorio 1998, Vorobyev *et al*. 1998). Human colour perception can be divided into two components: chromatic (hue and saturation) and achromatic (brightness) dimentions. These models are representations of the chromatic component of colour vision only (Renoult *et al*. 201).

Colour vision models require a minimum of four parameters for calculations: (1) photoreceptor sensitivity curves, (2) background reflectance spectrum, (3) illuminant spectrum, and (4) the observed object reflectance spectrum (stimulus). In addition, receptor noise limited models require photoreceptor noise for each photoreceptor type. Photoreceptor sensitivity curves are available for several animal taxa. If not available, the sensitivity curves can be estimated using formula based on wavelength at photoreceptor maximum sensitivity (*λ*_*max*_; Govardovskii *et al*. 2000). Background reflectance can be calculated by measuring the reflectance of materials found in the environment, such as leaves, twigs and tree bark. Alternatively, the background reflectance can be an achromatic spectrum of low reflectance value. The illuminant can be a reference spectrum (e.g. CIE standards), or, ideally, measured directly in the field using an irradiance measurement procedure (Endler 1990; 1993). Reflectance spectra are usually measured using a spectrometer (see Anderson & Prager 2006 for measurement procedures), but it can also be collected using photographic and hyperspectral cameras (Stevens *et al*. 2007; Chiao *et al*. 2011). All data must cover the same wavelength range as the photoreceptor sensitivity curves (300-700 nm for most cases).

I begin with Equations 1-5, which are common to all colour vision models presented here. Then calculation for each model is presented in a subtopic. I show formulae used to model trichromatic vision only. Formulae for tetrachromatic vision are available in the supplementary material. Photoreceptors are grouped by their maximum sensitivity value (*λ*_*max*_), from shortest to longest *λ*_*max*_. Honeybees workers (*Apis mellifera*), for instance, have three photoreceptor types with *λ*_*max*_ at ca. 344nm, 436nm and 544nm (Figure 1a; Peitsch *et al*. 1992).

**Figure 1.**
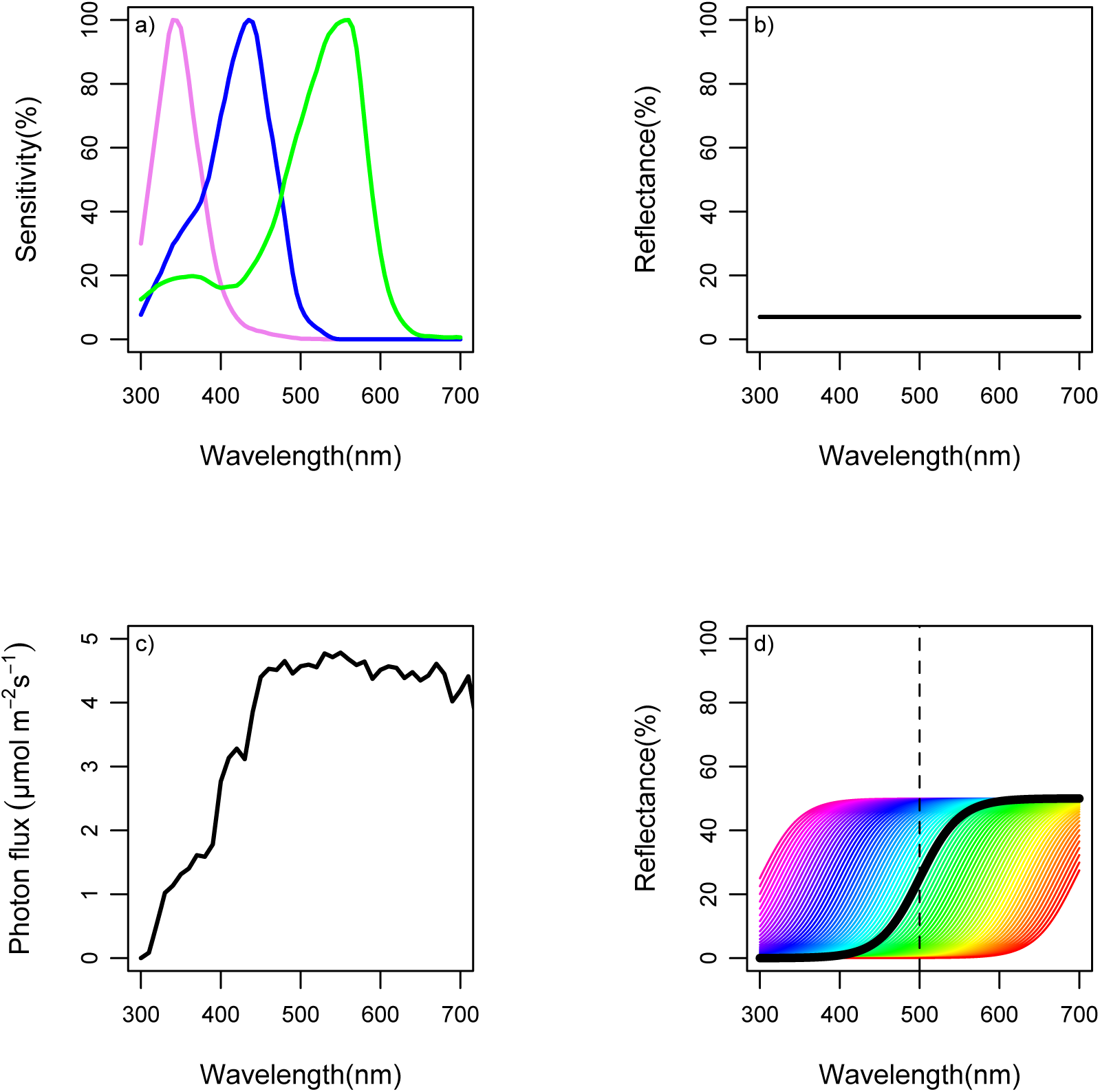
Basic setup used for colour vision model simulations. (a) Honeybee (*Apis mellifera*) photoreceptor sensitivity curves (data from Peitsch *et al*. 1992 available in Chittka & Kevan 2005); (b) Achromatic background reflectance spectrum; (c) CIE D65 standard daylight illuminant; and (d) Reflectance spectra generated by a logistic function with midpoints varying from 300 to 700nm at 5nm intervals. Spectrum colours are arbitrary. In black is shown a reflectance curve with midpoint at 500nm.

The first step is to calculate the total photon capture (*Q_i_*) of each photoreceptor type (*i*):

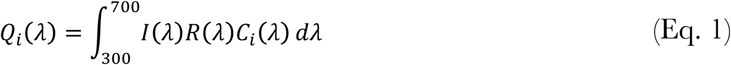
where *I* is the illuminant spectrum reaching the observed object, *R* is the reflectance of the observed object, *C_i_* is the photoreceptor sensitivity curve of photoreceptor *i*. The integration is usually done from 300 to 700nm, but this range can be changed depending on the animal of interest. Most mammals, for instance, do not capture photons below 400 nm. The second step is to calculate the photon catch by each photoreceptor (*i*) arising from the background reflectance:

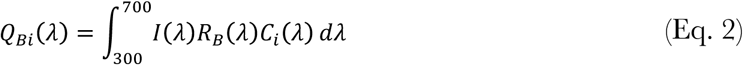

Where *I* and *C*_*i*_ are the same values in equation (1), and *R_B_* is the background reflectance. In practice photon catches are done by summation 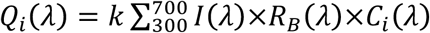, where *k* is the constant representing the interval between measurements, usually 1 nm. The relative photoreceptor photon catch (*q_i_*) is then calculated by:

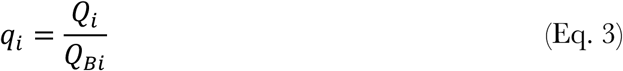

The rationale behind equation (3), referred as the von Kries transformation, is that photoreceptors are physiologically adapted to the light coming from the background, and that animals exhibit colour constancy (Chittka *et al*. 2014). So that if the environment is rich in wavelengths at the green region of the light spectrum, photoreceptors sensitive to this wavelength region will be less responsive.

### Colour hexagon model

The colour hexagon model (Chittka 1992) was formulated for hymenopteran vision. However, due its general form it can, and has been, applied for other taxa. Photoreceptor output (*E*) is given by:

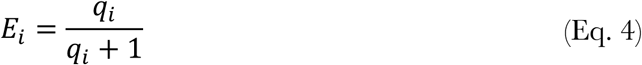

This means that photoreceptors output (*E*) will vary from 0 to 1, and its value will increase asymptotically to the limit of 1. This is done because the relationship between photoreceptor input-output is non-linear. E-values are then depicted into three vectors evenly distributed (120° between them). The resultant of receptor outputs is projected into a plan (chromaticity diagram) using the following formula:

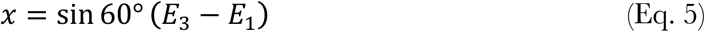

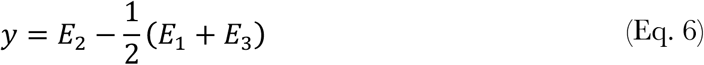

### Endler & Mielke (2005) model

The model is originally the first step for a statistical approach to study bird colouration as whole, not as individual colour patches (Endler & Mielke 2005). The model was adapted from tetrachromatic to trichromatic vision by Gomez (2006). The first step is to log-transform relative photon catches:

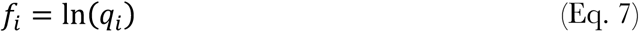

Then, *S*_*i*_ is transformed so that photoreceptor outputs *u* + *s* + *m* = 1:

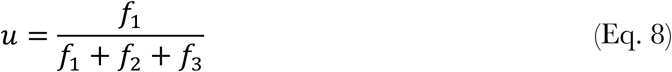

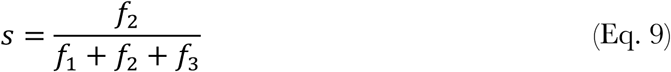

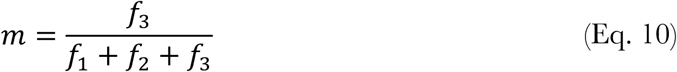

Rationale between equations 8-10 is that only the relative differences in photoreceptor outputs are used in a colour opponency mechanism. Photoreceptor outputs are projected into a triangular chromaticity diagram by the following formula (Gomez 2006):

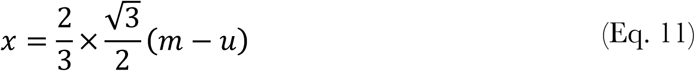

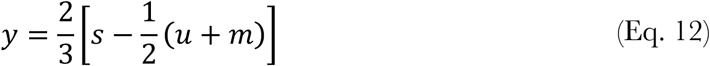

### Receptor noise limited models: linear and log-linear versions

The receptor noise limited model was developed to predict thresholds of colour vision. One of the assumption is that thresholds are given by noise arising at the receptor channels (Vorobyev & Osorio 1998). The first receptor noise limited model uses a linear relationship between photoreceptor input (*q_i_*) and output (*f_i_*) so that (linear version of the receptor noise limited model; Vorobyev & Osorio 1998):

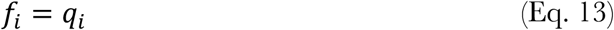

The log-linear version of receptor noise limited model assumes a log-linear relationship between photoreceptor input and output (log-linear version of the receptor noise limited model; (Vorobyev *et al*. 1998):

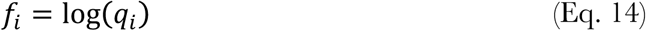

Equation (13) can be used when comparing colours that are very similar, otherwise equation (14) should be used. Then *f_i_* values are used to find the colour locus (*x, y*) in a chromaticity diagram (Hempel de Ibarra *et al*. 2014):

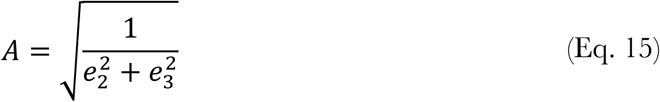

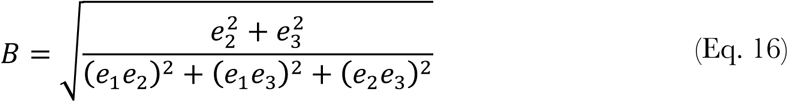

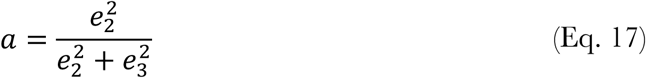

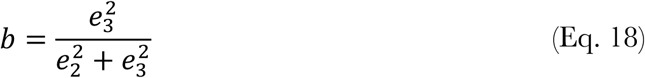

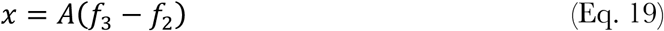

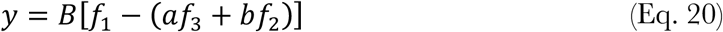

where *e*_*i*_ is the receptor noise of each photoreceptor, from shortest to longest wavelength. To date few species had their receptor noise (*e*_*i*_) measured directly (Vorobyev & Osorio 1998). In lack of a direct measurement, *e*_*i*_ can be estimated by the relative abundance of photoreceptor types in the retina and a measurement of a single photoreceptor noise-to-signal ratio:

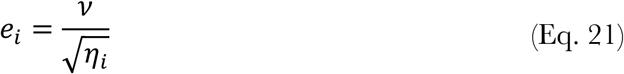

Where *v* is the noise-to-signal ratio of a single photoreceptor, and *η*_*i*_ is the relative abundance of photoreceptor *i* in the retina. Alternatively, *e*_*i*_ may be intensity depend (Renoult *et al*. 2017):

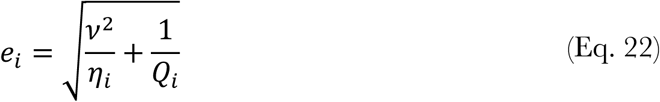
where *Q*_*i*_ is the photon catch given by equation (1). Equation (20) is usually valid in high light intensities, whereas equation (21) usually holds for dim light conditions (Vorobyev *et al*. 1998; Vorobyev & Osorio 1998).

### Distance between colour loci in chromaticity diagrams

Distances in chromaticity diagrams represent chromaticity similarities between two colours. The assumption is that the longest the distance, the more dissimilar two colours are perceived. Chromaticity distance between pair of reflectance spectra (*a* and *b*) are found by calculating the Euclidian distance between their colour loci (*x*, *y*) in the colour space:

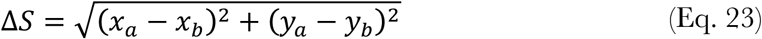

By definition, background reflectance lays at the centre of the background (*x* = 0, *y* = 0).

Therefore, the distance of the observed object against the background is given by:

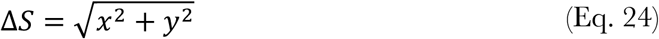

In the receptor noise models, Δ*S* between pair of reflectance spectra (*a* and *b*) can be calculated directly, without finding colour loci in the colour space (Vorobyev & Osorio 1998):

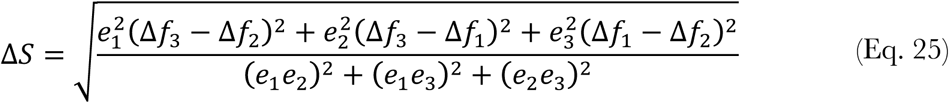

Where Δ*f_i_* is the difference between photoreceptor *i* output for the reflectance spectrum *a* and *b* (Δ*f_i_* = *f_ai_* − *f_bi_*). Using equation (24) will give the same value as calculating Δ*S* using equations (14-19) and then equation (23). In RNL models, Δ*S* = 1 equals one unit of just noticeable difference (JND). That means that, given the experimental conditions (large static object against a grey homogenous background), JND = 1 is the threshold for object detection; i.e. the minimum behaviourally discriminable difference between the object and the background.

## Simulations

I modelled the perception of the honeybee (*Apis mellifera*) using the colour vision models presented above: Chittka (1992) colour hexagon model (hereafter CH model), Endler & Milke (2005) model (hereafter EM model), and linear and log-linear versions of the receptor noise model (hereafter linear-RNL and log-RNL models (Vorobyev *et al*. 1998; Vorobyev & Osorio 1998). My aim was to compare between model results and analyse and illustrate how models behave in different scenarios. I begin with a basic model setup with simulated data. Then a make a series of changes to this basic model to investigate how models behave with typical input data used in ecology and evolution papers. At the end I use real flower reflectance data to compare model results.

### Simulation 01: Basic model setup

I used honeybee worker (*Apis mellifera*) photoreceptor sensitivity curves (data from Peitsch *et al*. 1992) available in Chittka & Kevan 2005); Figure 1a). As the background reflectance spectrum I created a theoretical achromatic reflectance with a constant 7% reflectance across 300 to 700nm (Figure 1b). As illuminant I used the CIE D65, a reference illuminant that correspond to midday open-air conditions (Figure 1c). For the receptor noise models I used measurements of honeybee photoreceptor noise (0.13, 0.06 and 0.12 for short, medium and long-wavelength photoreceptors; data from Peitsch 1992 available in Vorobyev & Brandt 1997). As the stimulus reflectance spectra I generated reflectance curves using a logistic function:

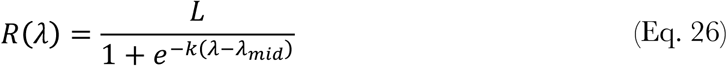

Where *R* is the reflectance value at wavelength *λ*, *L* gives the curve maximum reflectance value (%), *k* gives the steepness of the curve, and *λ_mid_* is the wavelength (nm) of midpoint. The logistic curve is a typical reflectance curve of many animal colour patches. I used a maximum value of *L*= 50% reflectance and a steepness of *k* = 0.04. I generated curves with midpoints varying from 300 to 700 nm with 5 nm intervals, in a total of 81 reflectance spectra (Figure 1d). For each model I calculated photoreceptor outputs, colour loci (*x* and *y*), and the chromatic distance to the background (Δ*S*) of each reflectance spectra using equations (1-24). In addition, as a supplementary material, I ran the same simulations with a tetrachromatic vision, and using a Gaussian function to generate the stimulus reflectance spectra (see Electronic Supplementary Material).

### Simulation 02: 10 percent point added to reflectance values

In the second simulation I added 10 percent point to the stimulus reflectance spectra (Figure 2a). My aim was to analyse how a relatively small change in reflectance curves affect model results. An increase in overall reflectance value can be an artefact of spectrometric measurement error (for guidance on spectrometric reflectance measurements see Anderson & Prager 2006).

**Figure 2.**
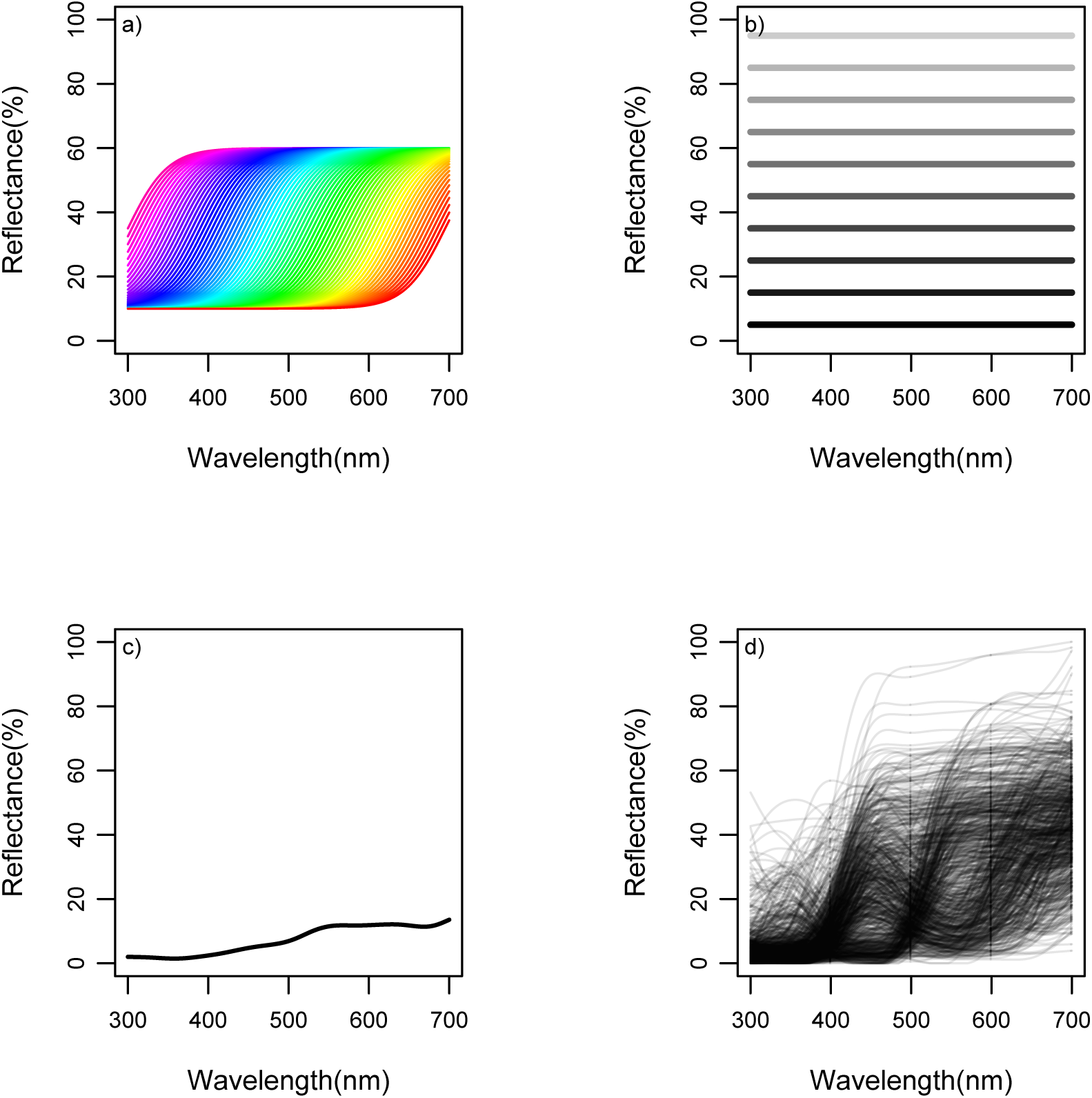
Changes from the basic setup used for colour vision model simulations. (a) Ten percent point added to the original reflectance spectra with midpoints varying from 300 to 700nm at 5nm intervals; (b) Achromatic reflectance spectra, with reflectance values from 5% to 95%, at 10 percent point intervals; (c) Background reflectance spectra calculated from the average reflectance of leafs, leaf litter, grasses and tree bark collected in the Brazilian savanna (data from Gawryszewski and Motta 2012); (c) Reflectance spectra of 859 flowers collected worldwide (data from the Flower Reflectance Database; Arnold *et al*. 2010).

### Simulation 03: achromatic reflectance spectra

Colour vision models are designed to deal with chromatic spectra (reflectance spectra that produces differences in photoreceptor outputs). However, some animal colours have reflectance spectra with a relatively constant reflectance value from 300 to 700nm, which we perceive as white, grey and black patches (achromatic variation). These type of spectra are sometimes modelled into colour vision models. In this simulation I generated a series of achromatic spectra with constant reflectance values from 300-700nm. I generated 10 reflectance spectra with reflectance values from 5% to 95%, with 10 percentage point intervals (Figure 2b).

### Simulation 04: Achromatic reflectance spectra and chromatic background reflectance spectrum

In the basic model I used an achromatic reflectance spectrum (7% reflectance from 300 to 700nm). In practice, however, most studies that apply colour vision models use chromatic reflectance backgrounds, such as leaf (e.g. Vorobyev *et al*. 1998), or an average of background material reflectance spectrum (e.g. Gawryszewski & Motta 2012). Models are constructed so that the background reflectance spectrum lie at the centre of the colour space. Vorobyev and Osorio (1998) specifically state that their linear receptor noise model is designed to predict perception of large targets, in bright light conditions and against an achromatic background. Despite of that, given that photoreceptors adapt to the light environment condition, usage of chromatic background is probably reasonable. Therefore, in this simulation I used the same achromatic reflectance spectra from simulation 03, but instead of having an achromatic background I used a chromatic background. The background is the average reflectance of leafs, leaf litter, tree bark and twigs collected in an area of savanna vegetation in Brazil (data from Gawryszewski & Motta 2012).

### Real reflectance data: comparison between models

In this setup my aim was to compare model results using real reflectance data. I used 858 reflectance spectra from flower parts collected worldwide and deposited in the Flower Reflectance Database (FReD; Arnold *et al*. 2010). I used only spectrum data that had a wavelength range from 300nm to 700nm. Data were then interpolated to 1nm intervals and negative values converted to zero. I used the same reflectance background from simulation 04, and other model parameters identical to the basic model setup. I compared models results visually, and by testing the pairwise correlation between model’s Δ*S* values. I used the Spearman correlation coefficient because data did not fulfil assumptions for a parametric test.

### Colourvision: R package for colour vision models and related functions

All calculation and figures presented here were performed using the colourvision R package. The package has functions for dichromatic, trichromatic and tetrachromatic linear and log-linear versions of the receptor noise limited model (Vorobyev *et al*. 1998; Vorobyev & Osorio 1998); and trichromatic and tetrachromatic versions of Chittka (1992) colour hexagon and Endler and Mielke (2005) models. Results from these models can be easily projected into their chromaticity diagrams for trichromatic and tetrachromatic vision. The colourvision package complements and can be used together with pavo R package for colour analyses (Maia *et al*. 2013), although it does not depend on it.

## Results

### Simulation 01: Basic model

Basic model results projected into chromaticity diagrams show differences between model predictions of colour perception for the same reflectance spectrum (Figure 3). CH model and the the linear-RNL model follow a similar path: data points follow a circular path that begins and ends near the centre of the colour diagram (Figure 3a and 3c). In the EM model, points follow two lines increasing in opposite directions, with data points reaching values outside colour space limits (Figure 3b). In the log-RNL model, points begin at the centre of the colour space and follow a curve increasing in distance from the centre of the colour space (Figure 3d).

**Figure 3.**
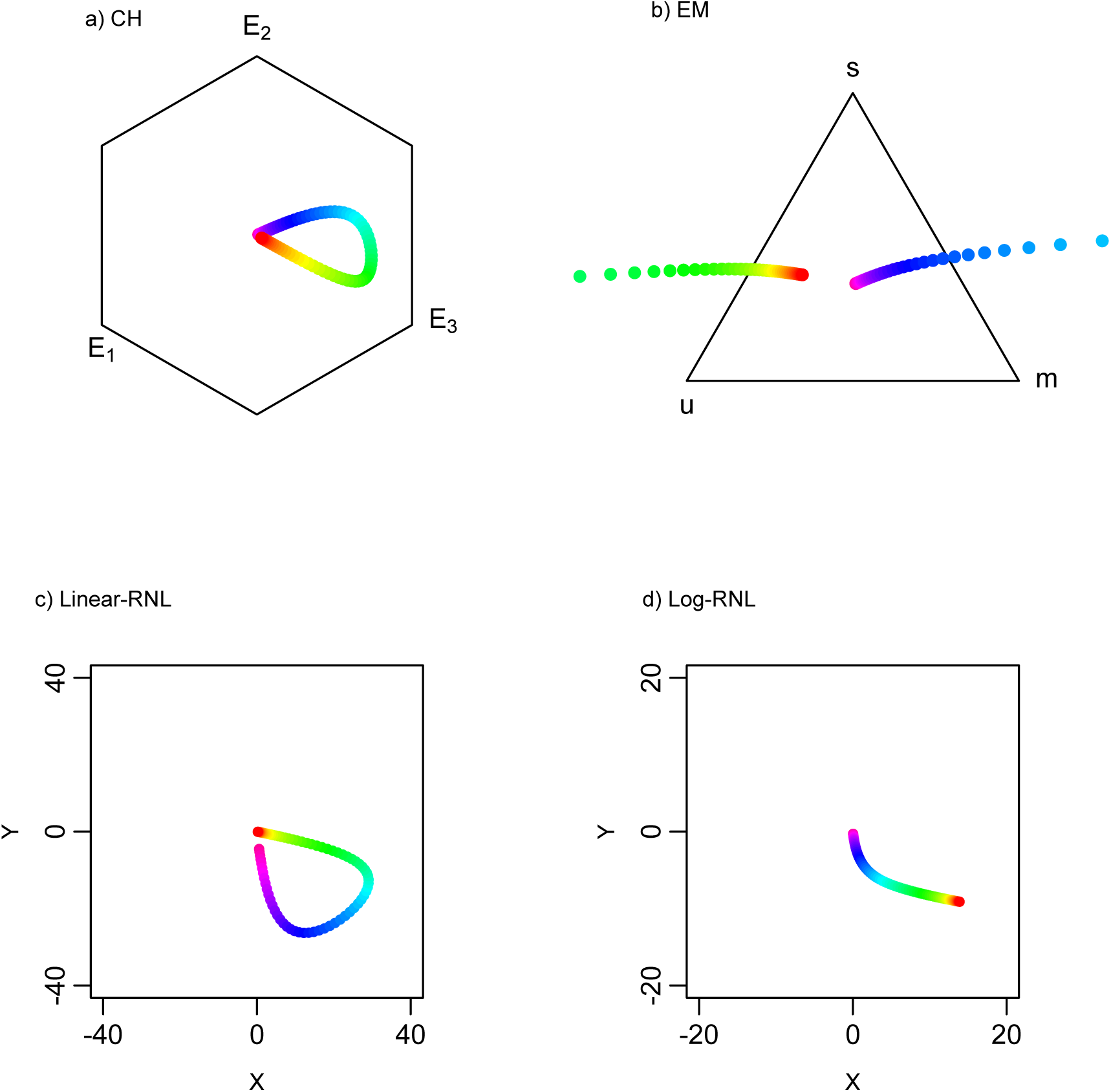
Chromaticity diagrams of the basic setup of colour vision model simulations: Chittka (1992) colour hexagon (CH), Endler & Mielke (2005) colour triangle (EM), and linear and log-linear Receptor Noise Limited models (Linear-RNL and Log-RNL; Vorobyev & Osorio 1998; Vorobyev *et al*. 1998). Colours correspond to reflectance spectra from Figure 1d.

CH model estimates a bell shaped Δ*S* curve against midpoint wavelength, with maximum Δ*S* for the reflectance curve with midpoint at 535nm (Figure 4a). Individual photoreceptors follow a sigmoid curve, with maximum values at short midpoint wavelengths and minimum values at long midpoint wavelengths (Figure 4a). EM model estimates unrealistic ΔS-values for reflectance curves with midpoints between 450-550nm (Figure 4b). A maximum Δ*S* = 116 is reached at 490nm midpoint wavelength (Figure 4b). Photoreceptor output also reach unrealistic negative values, and values above 1 (Figure 4b). This is consequence of equations 7-10: when *q_i_* is below 1, the ln-transformation generates negative values. Consequently, the denominator in equations 8-10 may reach values close to zero, which causes photoreceptor outputs to tend to infinity. Log-RNL model predicts a sigmoid ΔS curve, increasing from short to long midpoint wavelengths, reaching a maximum ΔS at at 700 nm (Figure 4d). Comparably to the EM model, the log-RNL model generates unrealistic negative photoreceptor excitation values (Figure 4d). Again, this happens because when *q_i_* is below 1 the log-transformation generates negative values (eq. 14). The linear-RNL version estimates a bell shaped ΔS curve, with a maximum Δ*S* at 470nm midpoint wavelength (Figure 4c). Photoreceptors present a sigmoid excitation curve, with maximum values at short midpoint wavelengths (Figure 4c).

**Figure 4.**
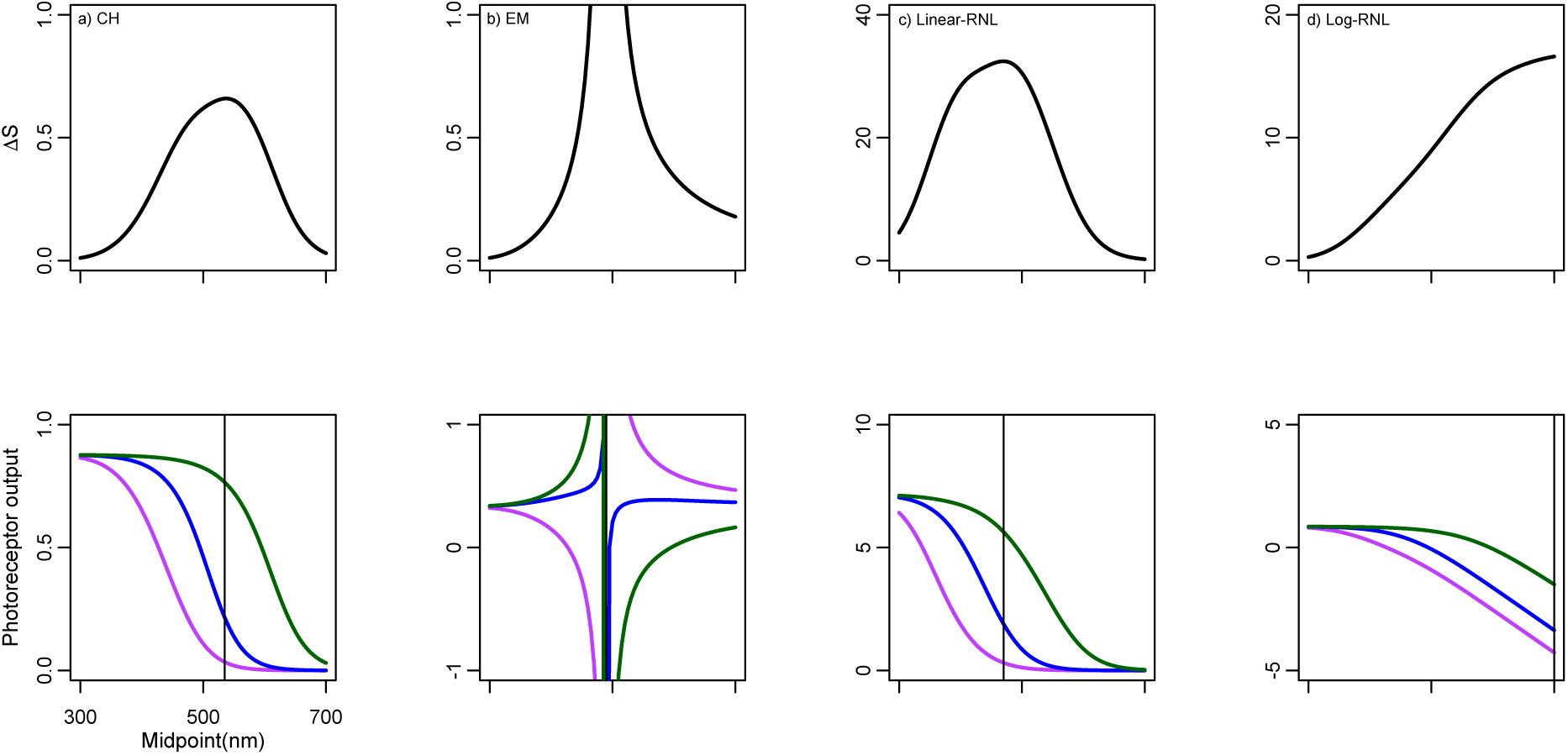
ΔS and photoreceptor outputs of the basic setup of colour vision model simulations (Figure 1): Chittka (1992) colour hexagon (CH), Endler & Mielke (2005) colour triangle (EM), and linear and log-linear Receptor Noise Limited models (Linear-RNL and Log-RNL; Vorobyev & Osorio 1998; Vorobyev *et al*. 1998). Variation in S-values as a function of reflectance spectra with midpoints from 300 to 700nm (top row). Photoreceptor output values as a function of the same reflectance spectra (bottom row). Violet, blue and green colours represent short, middle and long *λ*_max_ photoreceptor types. Vertical lines represent midpoint of maximum ΔS-values.

### Simulation 02: 10 percent point added to reflectance values

In this setup, models are more congruent in their results. Their chromaticity diagram indicates similar relative position of reflectance spectra between models (Figure 5). All of them estimate a bell shaped Δ*S* curve, with maximum values around 500 nm midpoint wavelength (Figure 6). CH model predicts a bell shaped ΔS curve with maximum Δ*S* peaking at 510nm (25 nm difference to the original model; Figure 6a). However, in comparison to the basic model there is an overall decrease in Δ*S* (Figure 3a and 6a). This happens because eq. 4 makes E-values non-linear as *q_i_* increases. Therefore, the relative differences between photoreceptors decreases and, as a consequence, Δ*S* decreases. Contrary to the basic model, EM model now estimates realistic ΔS and photoreceptor excitation values, with a peak at 540nm (Figure 6b). With a 10 percentage point increase in the reflectance, eq. 3 does not produce values below 1. As a consequence, eqs. 7-10 do not generate negative values and the denominator cannot reach near zero values. The same pattern occurs in the log-RNL model: no negative values are generated by eq. 14. Model estimates a bell shaped curve peaking at 505 nm (Figure 6d). The linear RNL model generates identical ΔS and colour loci values to the basic model. The 10 percentage point increase causes an increases in photoreceptor excitation values (Figure 6c). However, because the relative differences between photoreceptors remain the same, and the relationship between *q_i_* and *f* is linear (no transformation of *q_i_*) there is no difference in ΔS between the original and this model setup (Figures 4c and 6c).

**Figure 5.**
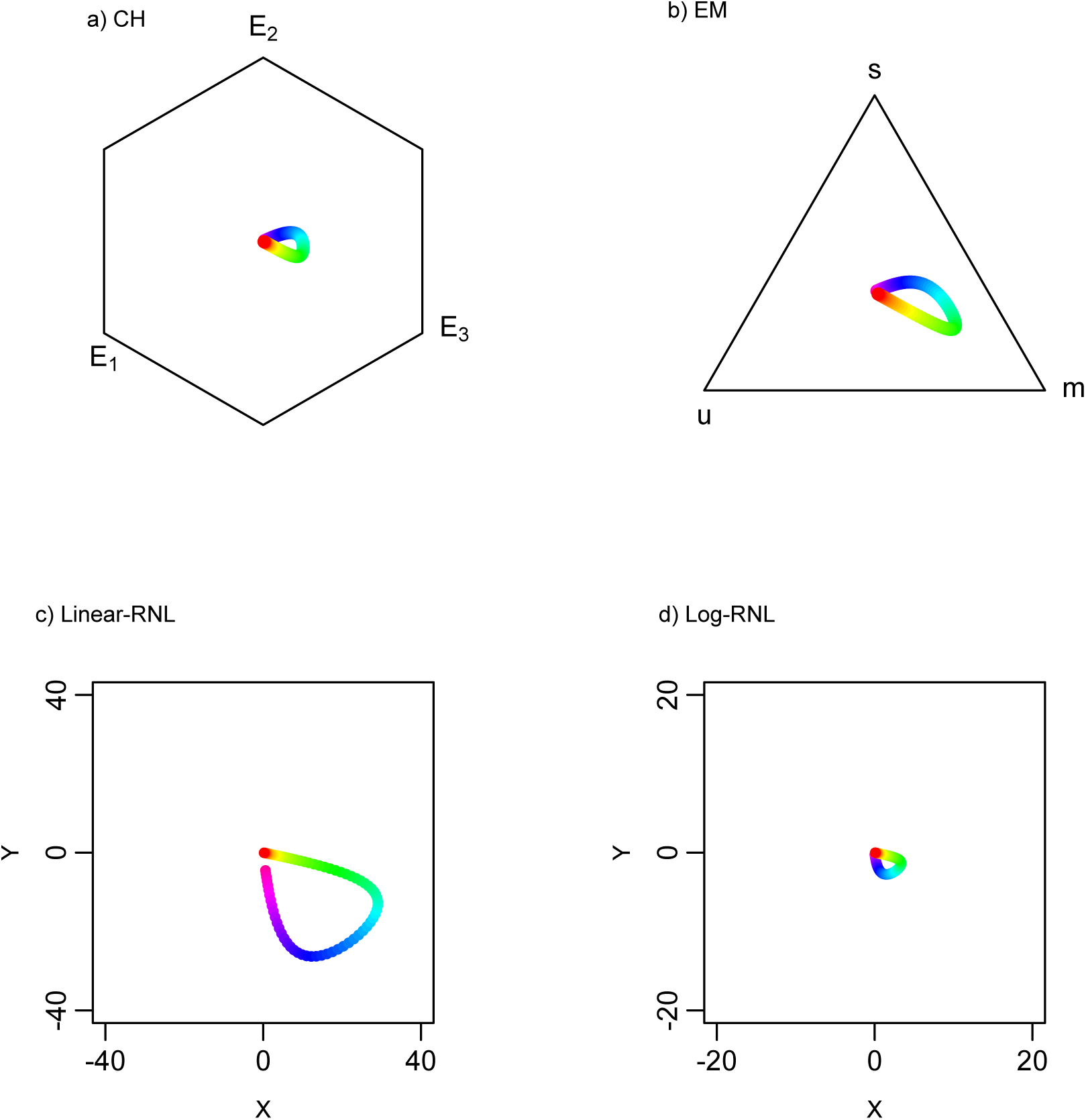
Chromaticity diagrams of the second simulation – 10 percent point added to reflectance values: Chittka (1992) colour hexagon (CH), Endler & Mielke (2005) colour triangle (EM), and linear and log-linear Receptor Noise Limited models (Linear-RNL and Log-RNL; Vorobyev & Osorio 1998; Vorobyev *et al*. 1998). Colours correspond to reflectance spectra from Figure 2a.

**Figure 6.**
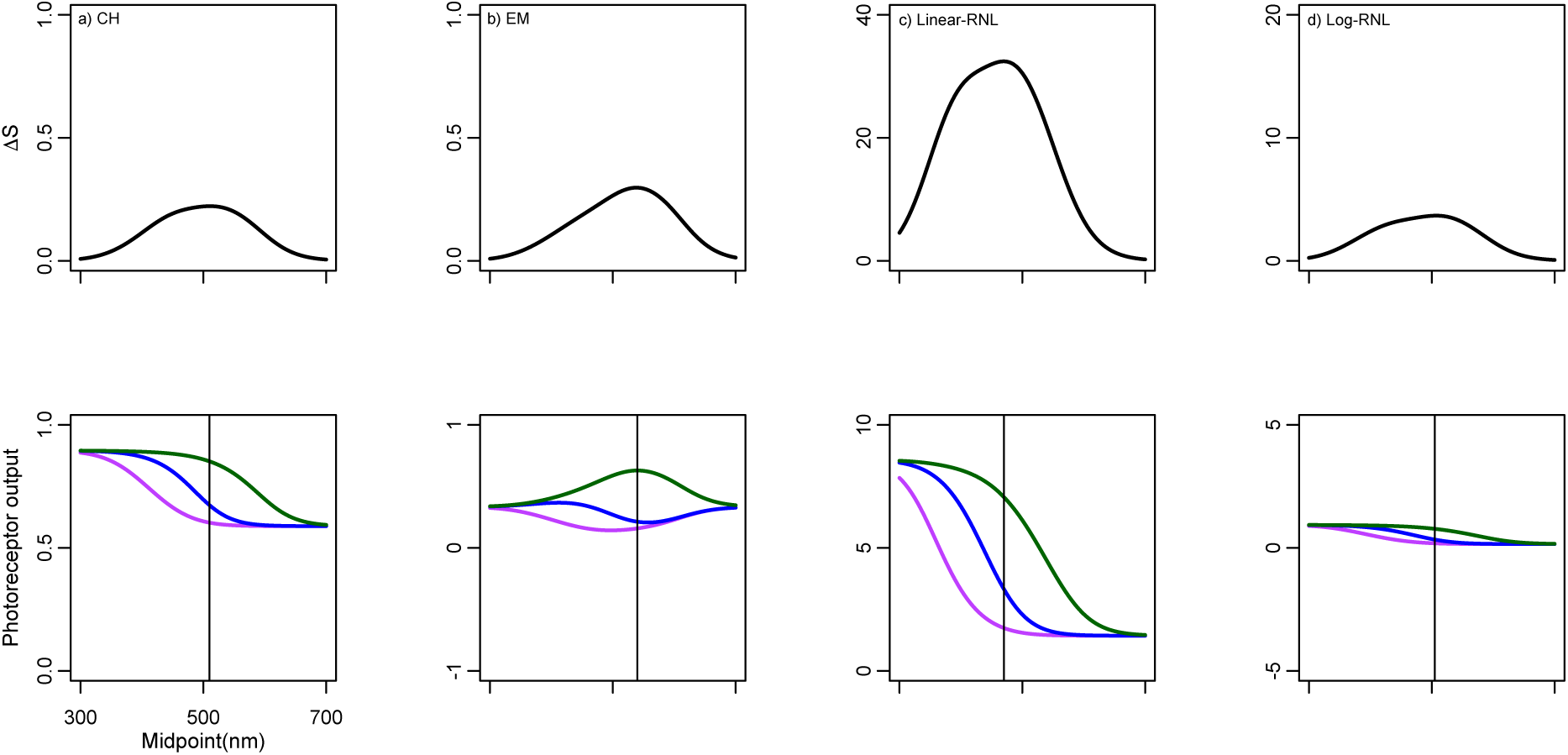
Δ*S* and photoreceptor outputs of the second setup of colour vision model simulations - 10 percent point added to stimulus reflectance spectra: Chittka (1992) colour hexagon (CH), Endler & Mielke (2005) colour triangle (EM), and linear and log-linear Receptor Noise Limited models (Linear-RNL and Log-RNL; Vorobyev & Osorio 1998; Vorobyev *et al*. 1998). Variation in ΔS-values as a function of reflectance spectra with midpoints from 300 to 700nm (top row). Photoreceptor output values as a function of the same reflectance spectra (bottom row). Violet, blue and green colours represent short, middle and long *λ*_max_ photoreceptor types. Vertical lines represent midpoint of maximum ΔS-values. For comparison, scales are the same as in Figure 4.

### Simulation 03: Achromatic reflectance spectra

With achromatic reflectance spectra all datapoints are in the centre of the colour diagram. Consequently, ΔS for all models and all reflectance values equals zero. This happens because all three photoreceptors respond equally to the achromatic reflectance spectra. Nonetheless, the type of response varies between models. In the CH model, photoreceptor output increases asymptotically as reflectance increases, which is result of eq. 4. In the EM model, photoreceptor outputs are not affected by variation in reflectance values. This happens because EM model considers only the relative differences between photoreceptors response (eqs. 8-10). In the RNL models, photoreceptor output increases linearly in the linear version, and asymptotically in the log version.

### Simulation 04: Achromatic reflectance spectra and chromatic background reflectance spectrum

Model results of achromatic reflectance spectra against a chromatic background differ to the model predictions when the background is achromatic (Figure 7 and Figure 8). The chromatic background causes differences in photoreceptor outputs. Consequently, achromatic reflectance spectra do not lay at the centre of the colour spaces. The CH model shows a maximum ΔS values of 0.31 at 5% reflectance achromatic spectrum (Figure 8a). ΔS values then decrease as the reflectance value of achromatic spectra increases (Figure 8a). Photoreceptor output values converge to the asymptote as the reflectance value of achromatic spectra increases (Figure 8a). EM model produce spurious values at 5% reflectance achromatic spectrum because it generates negative photoreceptor output values (Figure 8b). From 15% beyond, ΔS values then decrease as the reflectance value of achromatic spectra increases (Figure 8b). The linear-RNL model shows a linear increase in ΔS values as the reflectance value of achromatic spectra increases (Figure 8c). Similarly, photoreceptor outputs also increase linearly as as the reflectance value of achromatic spectra increases, but with different slopes for each photoreceptor type (Figure 8c). Contrary to other models, ΔS-values in the log-RNL model do not change with varying reflectance value of achromatic spectra (Figure 8d). Although photoreceptor outputs increase as reflectance value of achromatic spectra increases (Figure 8d), the difference between photoreceptor outputs remains the same. Consequently, ΔS-values do not change.

**Figure 7.**
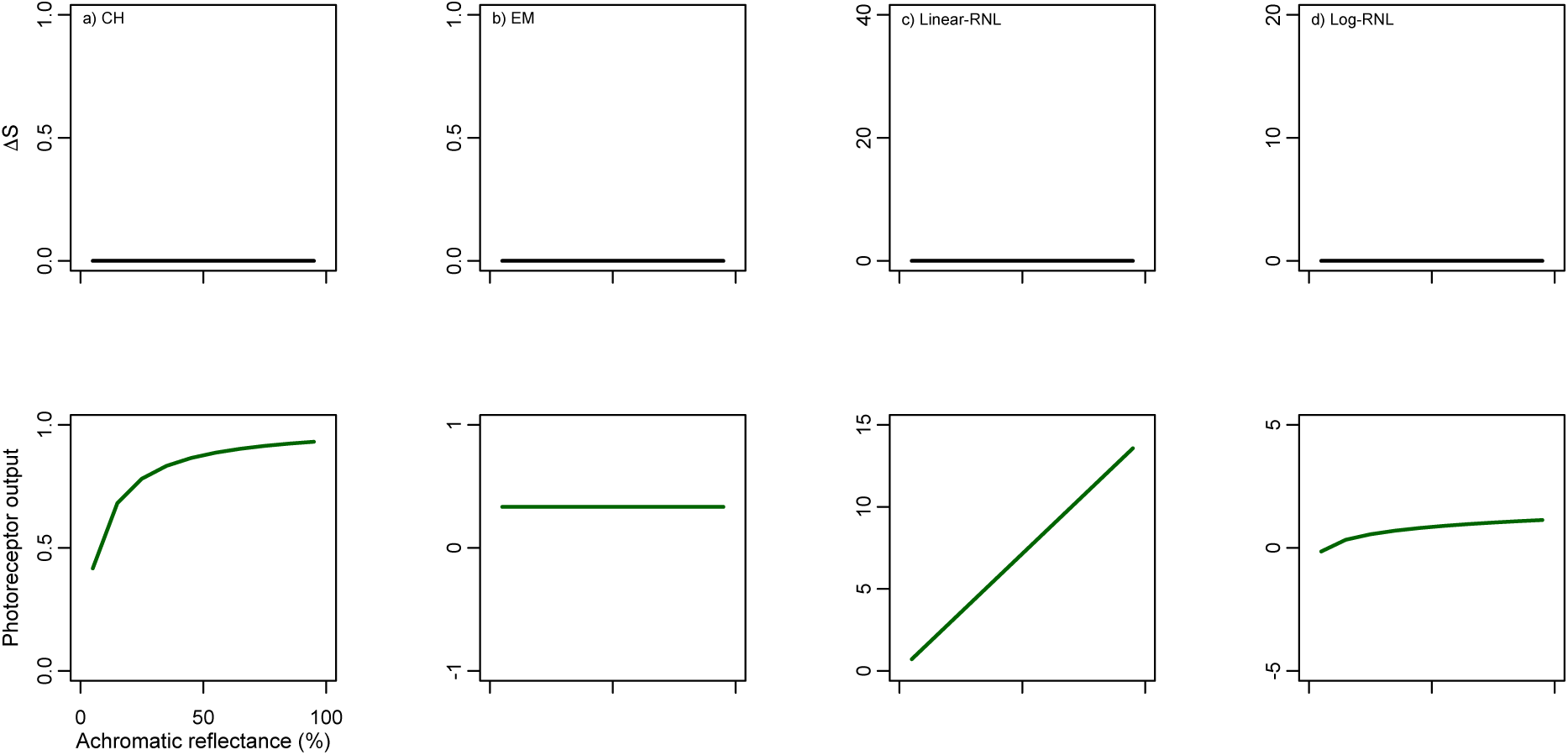
ΔS and photoreceptor outputs of the third setup of colour vision model simulations – achromatic stimulus against achromatic background: Chittka (1992) colour hexagon (CH), Endler & Mielke (2005) colour triangle (EM), and linear and log-linear Receptor Noise Limited models (Linear-RNL and Log-RNL; Vorobyev & Osorio 1998; Vorobyev *et al*. 1998). Variation in ΔS-values as a function of spectra with achromatic reflectance from 5% to 95% (top row). Photoreceptor output values as a function of the same reflectance spectra (bottom row). Photoreceptors are colour coded by their *λ*_max_ photoreceptor, however they do not appear because are all superimposed. With the exception of c) Linear-RNL, scales are the same as in Figure 4.

**Figure 8.**
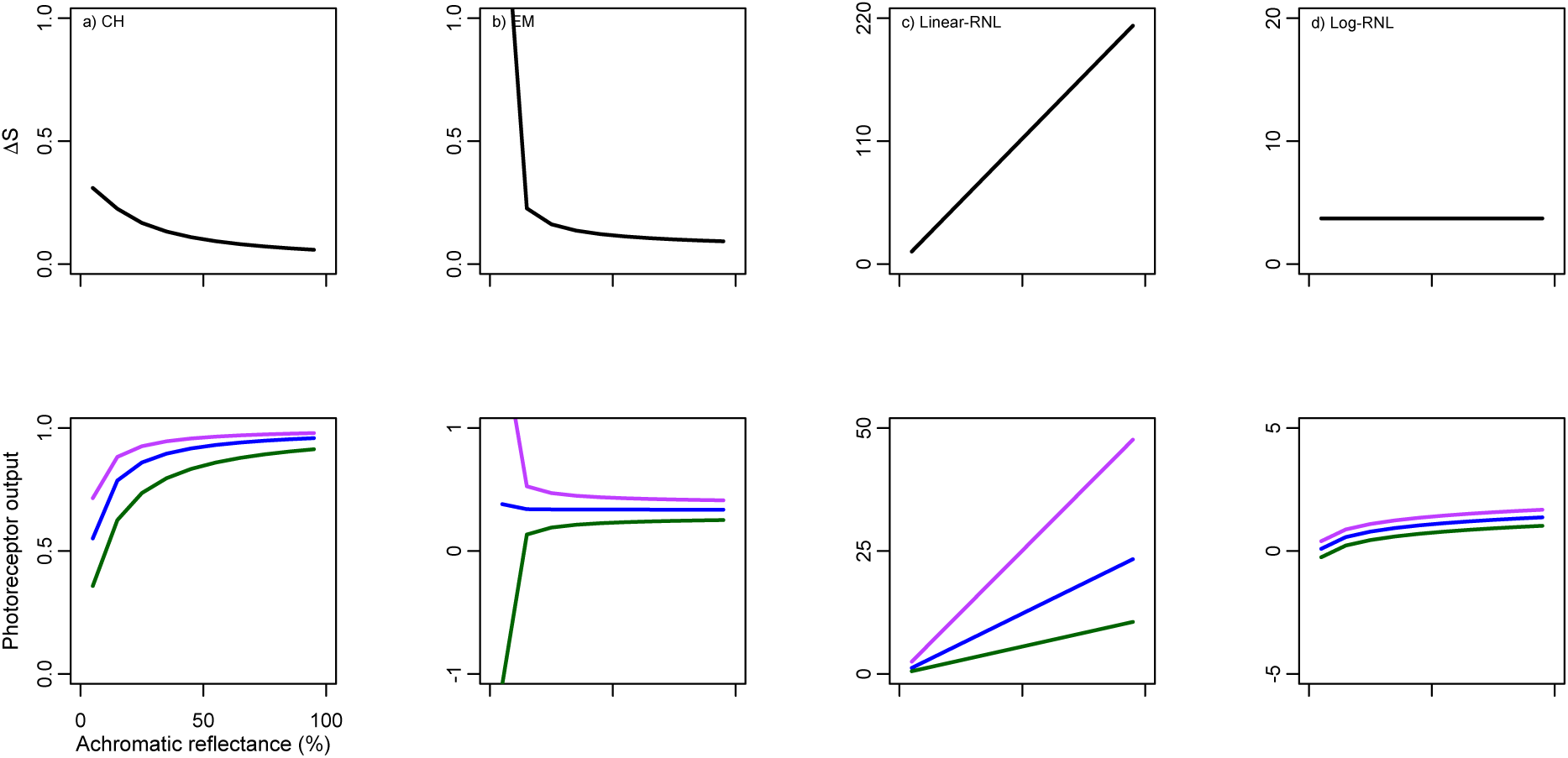
ΔS and photoreceptor outputs of the fourth setup of colour vision model simulations – achromatic stimulus against chromatic background: Chittka (1992) colour hexagon (CH), Endler & Mielke (2005) colour triangle (EM), and linear and log-linear Receptor Noise Limited models (Linear-RNL and Log-RNL; Vorobyev & Osorio 1998; Vorobyev *et al*. 1998). Variation in ΔS-values as a function of spectra with achromatic reflectance from 5% to 95% (top row). Photoreceptor output values as a function of the same reflectance spectra (bottom row). Violet, blue and green colours represent short, middle and long *λ*_max_ photoreceptor types. With the exception of c) Linear-RNL, scales are the same as in Figure 4.

### Simulation 05: Real reflectance data

When real flower reflectance spectra are used, models also give different relative perception for the same reflectance spectrum. The results of the CH model and the log-RNL model are similar both qualitatively and quantitatively: colour loci projected into the colour space (Figure 9) show similar relative position of reflectance spectra; and there is a high correlation score between ΔS values (Figures 9a and 9d; ρ=0.884; N=858; S=12165000; p<0.001). Even though many EM points lay outside the chromaticity, results suggest a high agreement between CH and EM models (Figure 9a and 9b; ρ=0.889; N=858; S=11718000; p<0.001). There was a moderate agreement between the linear and log version of the RNL model (Figures 9c and 9d; ρ=0.434; N=858; S=59623000; p<0.001, P<0.001), and between EM and log-RNL models (Figure 9b and 9d; ρ=0.662; N=858; S=35572000; p<0.001). There was a poor agreement between the linear-RNL and both EM models (Figure 9b and 9c; (ρ=-0.264; N=858; S=133060000; p<0.001), and CH models (Figure 9a and 9c; ρ=0.037; N=858; S=101370000; p=0.278)

**Figure 9.**
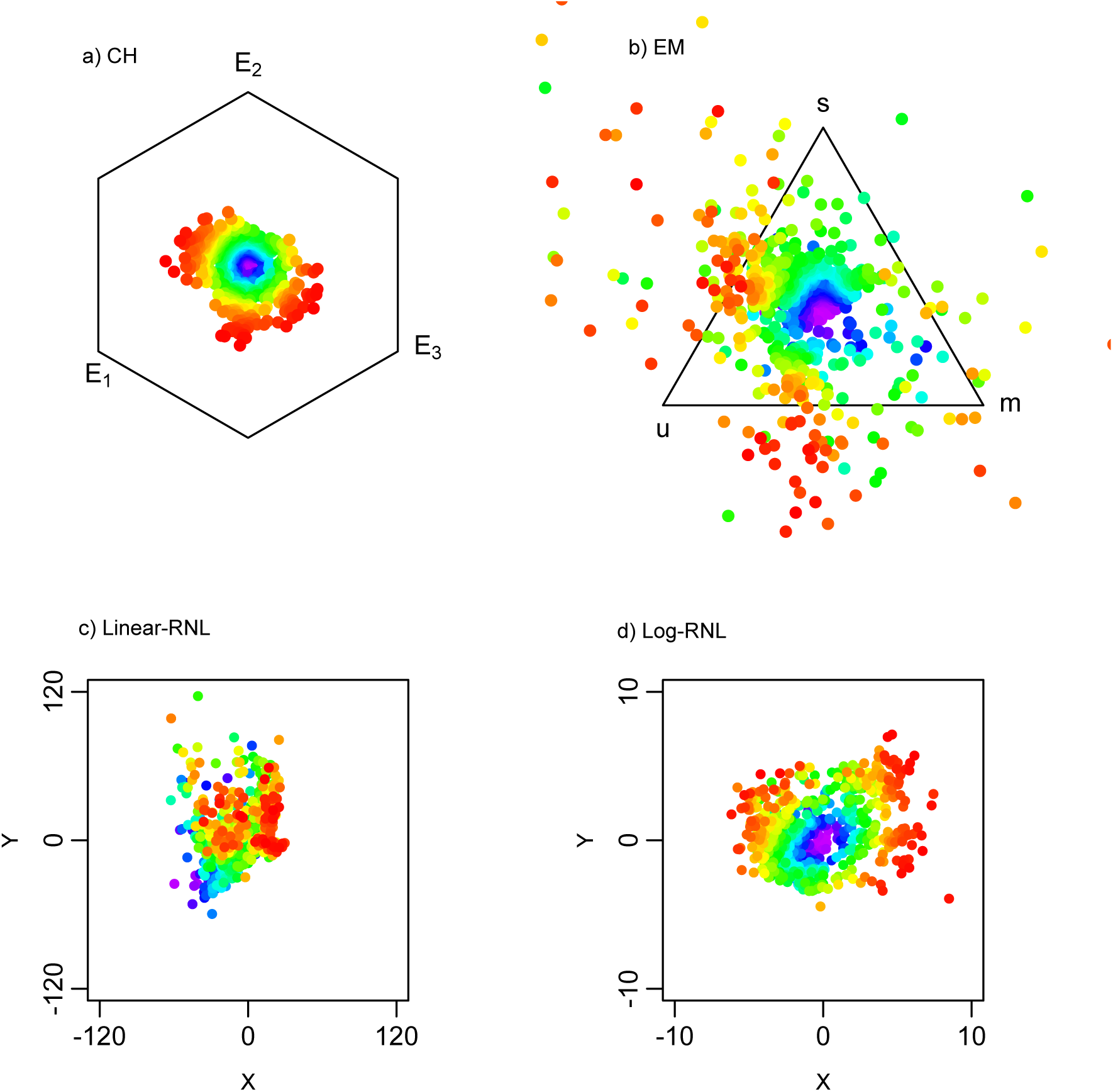
Flower reflectance spectra (N=858) projected into chromaticity diagrams: Chittka (1992) colour hexagon (CH), Endler & Mielke (2005) colour triangle (EM), and linear and log-linear Receptor Noise Limited models (Linear-RNL and Log-RNL; Vorobyev & Osorio 1998; Vorobyev *et al*. 1998). To facilitate model comparison, point colours correspond to chromaticity distances in the CH chromaticity diagram.

## Discussion

Application of colour vision models are now widespread in several fields of ecology and evolutionary biology. However, simulations presented here show that under certain conditions these models do not agree, and can produce spurious results. As any model, colour vision models have been developed based on a set of assumptions. Knowledge of model strength and limitation are crucial to the correct application and interpretation of colour vision model results.

Colour vision models, in special models that are log-transformed (eqs. 7 and 14) and convert photoreceptor output to relative values (eqs. 8-10) are prone to produce unrealistic results when the observed reflectance generates a lower response than the background (i.e. *Q_*i*_* < *Q_*Bi*_*). The log-transformation (and the transformation in the CH model) is behaviourally justified due to the Weber–Fechner law of psychophysics (Renoult *et al*. 2017). The law states that the perceived difference between a pair of stimuli has a non-linear relationship with their absolute difference. I.e., humans perceive 150g and 100g weights as more dissimilar than 1150g and 1100g weights. This is illustrated by the photoreceptor outputs in Figure 7. In this simulation the achromatic reflectance spectrum increases from 10% to 90% by 10 percept point steps. In the linear-RNL model, photoreceptor outputs respond linearly to the 10% increase, so that the difference in receptor outputs is the same between the 10% vs 20% reflectance spectra as between the 80% vs 90% reflectance spectra. In the log-RNL model, however, the difference between 10% vs. 20% is greater than the difference between 80% vs 90%.

In addition, low reflectance achromatic spectra (i.e. dark colour patches) may also produce spurious values when the background is chromatic (Figure 8), because at small reflectance values, small differences between photoreceptor outputs may be large in proportion to photoreceptor outputs, and consequently generate large Δ*S* values. Interpretation of model results depends on detailed knowledge of how models are calculated. Inspection of individual photoreceptor outputs can give insights into colour loci (*x*, *y*) and chromaticity distance (Δ*S*) values.

Comprehension of the physiology of vision of the animal observing the scene is also imperative. Honeybees, for instance, use colour vision only when the observed object subtend a visual angle larger than ca. 15° (Giurfa *et al*. 1996). Moreover, bees appear to completely ignore brightness when using the chromatic channel (Giurfa *et al*. 1997), so that equation (20) holds even in low light conditions (Vorobyev & Osorio 1998). In humans, on the other hand, the achromatic dimension appears to dominate in dim light conditions (King-Smith & Carden 1976; Vorobyev & Osorio 1998). These models also do not incorporate higher order cognition abilities that may affect how colour are perceived (Dyer 2012). In bees, for instance, previous experience, learning and experimental conditions may affect their behavioural discriminability thresholds (Chittka *et al*. 2003; Dyer & Chittka 2004; Giurfa 2004; Dyer *et al*. 2011; Dyer 2012); and in humans the ability to discriminate between colours is affected by the existence of linguistic differences for colours (Winawer *et al*. 2007).

A common misconception arises from the use of detectability/discriminability thresholds. The RNL model for instance, predicts well the detectability of monochromatic light against a grey background. For this model, and given the experimental condition, a Δ*S* = 1 equals one unit of just noticeable difference (JND; Vorobyev & Osorio 1998). Stimuli with values equal or above 1 can be detectable against the background, under the experimental condition. However, this threshold is not fixed. It can vary depending on the background, on the chromatic difference between the object and the background, and on the subject previous experience. For zebra finches, for instance, the same pair of similar red object have a discriminability threshold of ca. 1 JND when the background is red, but much higher when the background is green (Lind 2016). The same study emphasises the difference between detecting one object against the background and discriminating two similar objects: detection thresholds are usually higher than discrimination thresholds (as measured by the RNL model; Lind 2016). Given the variation in thresholds, it is misleading to interpret Δ*S* values as binary variable: i.e. above the threshold, detectable; below threshold, not detectable. Instead, use of Δ*S* values as they are, a continuum, makes the interpretation more realistic. E.g., a stimulus with JND = 2 is likely chromatically similar to the background, and is possibly more often not detected than a stimulus with JND = 5.

In addition, models presented here are pairwise comparison between colour patches, which do not incorporate the complexity of an animal colour pattern composed by a mosaic of colour patches of variable sizes. Endler and Mielke (2005) provide a methodological and statistical tool that can deal with a cloud of points representing an organism colour patches. Use of hyperspectral cameras or adapted DSLR cameras may facilitate the analysis of animal colouration as a whole (Stevens *et al*. 2007; Chiao *et al*. 2011). Other aspects that may be important when detecting a target, such as size, movement, and light polarization (Cronin *et al*. 2014), are also not incorporated into those models.

In conclusion, colour vision models are extremely useful and can provide insightful results on ecological and evolutionary aspects of colour in nature. Nonetheless, they should be regarded as an approximation of the perceived differences between pairs colours by a particular organism. Good application of colour vision models depends on the inspection of photoreceptor output values, knowledge of model assumptions, comprehension of the mathematical formula behind each model and familiarity with mechanisms of colour vision of the animal being modelled. Comparison of model results with field and laboratory based behavioural experiments are also crucial to complement and validate model results.

